# Deuterium-reinforced polyunsaturated fatty acids protect against muscle atrophy induced by type 1 diabetes in mice

**DOI:** 10.1101/2024.12.17.627433

**Authors:** Hiroaki Eshima, Ishihara Tomoaki, Ayaka Tabuchi, Yutaka Kano, Kenji Kurokawa, Mikhail S. Shchepinov

**Author notes:** Correspondence to: Hiroaki Eshima, Ph.D.,; Mikhail S. Shchepinov, Ph.D.

## Abstract

**HIGHLIGHTS:**

- D-PUFA diet prevents muscle atrophy in STZ-induced diabetic mice.
- D-PUFA diet prevents muscle weakness depending on increased calcium release in STZ-induced diabetic mice.
- D-PUFA diet may show a trend to decrease blood glucose in STZ-induced diabetic mice.
- D-PUFA diet does not alter ferroptosis-related protein profiles including ACSL4, LPCAT3, ALOX12, and Gpx4.

Oxidative stress and reactive oxygen species (ROS) have been linked to muscle atrophy and weakness. Diabetes increases the oxidative status of lipoproteins in nearly all tissues, including muscle tissues, but the role of lipid ROS on diabetes-induced muscle atrophy is not fully understood. Deuterium reinforced polyunsaturated fatty acids (D-PUFA) are more resistant to ROS-initiated chain reaction of lipid peroxidation than regular hydrogenated PUFA (H-PUFA). In this study, we tested the hypothesis that D-PUFA would protect muscle atrophy induced by diabetes driven by an accumulation of lipid hydroperoxides (LOOH). C57BL/6J mice were dosed with H-PUFA or D-PUFA for four weeks through dietary supplementation and then injected with streptozotocin (STZ) to induce insulin-deficient diabetes. After two weeks, muscles tissues were analyzed for individual muscle mass, force generating capacity and cross-sectional area. Skeletal muscle fibers from diabetic mice exhibited increased total ROS and LOOH. This was abolished by the D-PUFA supplementation regardless of accumulated iron. D-PUFA were found to be protective against muscle atrophy and weakness from STZ-induced diabetes. Prevention of muscle atrophy and weakness by D-PUFA might be independent of ACSL4/LPCAT3/15-LOX pathway. These findings provide novel insights into the role of LOOH in the mechanistic link between oxidative stress and diabetic myopathy and suggest a novel therapeutic approach to diabetes-associated muscle weakness.

## 1. Introduction

Diabetes represents a major public health concern that has a considerable impact on healthcare expenditures. It also induces muscle atrophy and weakness leading to limited functional capacity and quality of life [1, 2]. Muscle atrophy induced by diabetes may be associated with lower protein synthesis rates and increased protein breakdown [3, 4]; however, the upstream mechanism by which these changes occur is not well understood.

Reactive oxygen (ROS) and reactive carbonyl species (RCS) are suspected to be linked to the skeletal muscle atrophy [5, 6]. We and others demonstrated the association of elevated levels of reactive lipid carbonyls with muscle atrophy and weakness following inactivity [7, 8] and sarcopenia [9, 10]. These dysfunctions are mitigated by preventing the accumulation of reactive lipid carbonyls with interventions such as overexpression of glutathione peroxidase 4 (Gpx4) [8, 11] and inhibition of lysophosphatidylcholine acyltransferase 3 (LPCAT3) [7]. A recent study found that high glucose accumulates reactive lipid carbonyls and causes cell death in C2C12 cell line, suggesting that hyperglycemic condition may be associated with reactive lipid carbonyls [12]. Moreover, empagliflozin, a clinical hypoglycemic gliflozin drug, inhibits reactive lipid carbonyls and enhances muscle cell survival by restoring the expression of Gpx4. Taken together, reactive lipid carbonyls may contribute to muscle atrophy induced by diabetes; while the effect of inhibiting reactive lipid carbonyls on diabetes-induced muscle atrophy and weakness has not yet been investigated.

Lipid peroxidation is initiated by prooxidants such as hydroxyl radicals attacking the carbon-carbon double bond in fatty acids, particularly the polyunsaturated fatty acids (PUFA) - containing phospholipids [13]. By contrast, deuterium atoms incorporated into PUFA at bis-allylic positions (D-PUFA) give rise to kinetic isotope effect (KIE), slowing down the rate-limiting step of the chain process. The effect is further amplified along the chain, essentially resulting in the inhibition of lipid peroxidation. Previous studies demonstrated that dietary D-PUFA partially protect against cellular damage by Parkinson’s disease and atherosclerosis by lowering lipid peroxidation [14–17]. However, it is unknown if D-PUFA is sufficient to prevent muscle atrophy and weakness induced by diabetes. Thus, the purpose of this study was to test the hypothesis that D-PUFA would protect an accumulation of lipid peroxidation, thereby muscle atrophy and weakness induced by diabetes.

## 2. Materials and methods

### 2.1. Animals

Forty C57BL/6J mice were purchased from Nippon SLC (5 weeks old, Hamamatsu). The mice were maintained under a 12:12-h light-dark cycle with ad libitum access to food and water, except when they were fasted. The procedures used were conducted by the Guide for the Care and Use of Laboratory Animals as adopted and promulgated by the Declaration of Helsinki and the Faculty of Pharmaceutical Science using the Nagasaki International University Publication, which was enacted in 2009. All experiments were approved by the Animal Experimental Committee of Nagasaki International University (approve No. 157 and 170).

### 2.2. PUFA diets

After a 1 wk initial acclimatization period, mice were randomly fed an AIN-93M-based diet (Research Diets, USA) containing either control PUFAs (H-PUFA) or the isotope-reinforced PUFAs (D-PUFA). The composition of all diets is presented as previously described [17]. Food intake was monitored per cage once per week. After 4 wks of feeding, mice were given an injection of streptozotocin (STZ). Following the injection of STZ, mice were fed continued H-PUFA or D-PUFA for 2 weeks (Fig. 1A).

**Figure 1.**
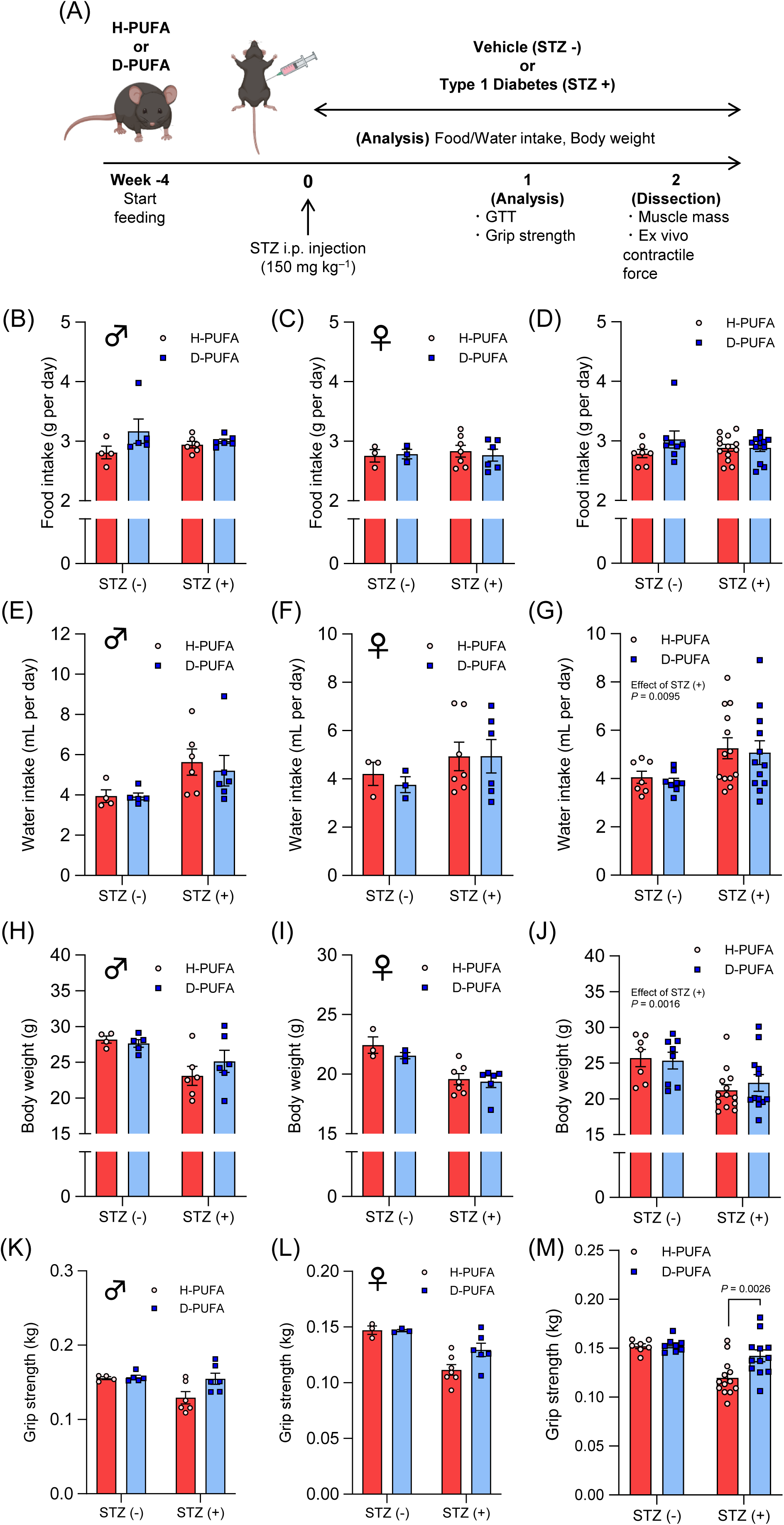
Physiological parameters in Vehicle (STZ -) and Type 1 diabetic (STZ +) mice fed H-PUFA and or D-PUFA. (A) Schematic of study design. (B-D) Food intake. (B) Male (♂) (C) Female (♀) and (D) Total mice (n = 7-13/group). (E-G) Water intake. (E) Male (♂) (F) Female (♀) and (G) Total mice (n = 7-13/group). (H-J) Body weight. (H) Male (♂) (I) Female (♀) and (G) Total mice (n = 7-13/group). (K-M) Grip strength. (K) Male (♂) (L) Female (♀) and (M) Total mice (n = 7-13/group). For the group comparison, two-way ANOVA were performed with Sidak’s post-hoc test. All data are mean ± SEM.

### 2.3. STZ-induced diabetic model

Mice were given an intraperitoneal injection of 150 mg/kg body wt of STZ (S0130, Sigma-Aldrich St. Louis, MO) prepared fresh in 0.1 M citrate buffer (pH 4.5) or injected with an equivalent volume of vehicle as a control. Mice were monitored for hyperglycaemia on day 3 using a One Touch glucometer (Lifescan, Johnson & Johnson, Switzerland), and only mice in which random blood glucose levels reached >300 mg dl^-1^ were used for experiments. Mice were euthanized on day 14 following STZ injection.

### 2.4. Glucose tolerance test

The intraperitoneal glucose tolerance test (GTT; 16–18 h; 0.5 g/kg body wt) was performed, as previously described [18, 19]. Blood samples were collected from the tail of mice at 0, 15-, 30-, 60-, and 120-min. Blood glucose concentrations were measured using a One Touch glucometer.

### 2.5. Limb grip strength test

The limb grip strength test was performed, as previously described [20]. The mice grasping the bar were pulled vertically by an inspector, and peak pull force was recorded on a digital force transducer (GPM-100; Melquest, Toyama, Japan). Tests were performed as 6 consecutive measurements per day at one-minute intervals. The average of 6 measurements was used to represent grip strength for each mouse. All tests were performed on the light cycle in the room where the animals were housed.

### 2.6. Tissue preparation

Mice were anesthetized via isoflurane (Mylan Pharmaceuticals, Osaka, Japan), and their muscles were dissected once a surgical level of anesthesia was reached. The tibialis anterior (TA), extensor digitorum longus (EDL), soleus (SOL), gastrocnemius (GAS), and Quadriceps (QUA) were carefully dissected from mice and quickly weighed before either being assessed for ex vivo force production, flash frozen in liquid nitrogen, or preserved in optimal cutting temperature (OCT) compound. The flexor digitorum brevis (FDB) muscles were dissected from mice for intracellular Ca^2+^ measurement. The abdominal visceral fat, epididymal fat, and liver were collected and weighed [19].

### 2.7. Ex vivo skeletal muscle force production

The force-frequency relationship was assessed in intact EDL muscles, as previously described [18, 19, 21–23]. Briefly, isolated EDL preparations were mounted on a force transducer (UL-100; Minebea, Tokyo, Japan), and the hock was fixed in a chamber containing oxygenated Krebs-Henseleit Buffer solution at 30°C. The optimal length of the muscle was determined via maximum twitch force production. Then, the isolated muscle was stimulated with 500-ms trains of current pulses at 10, 20, 40, 60, 80, 100, 125, 150, and 200 Hz at 1-min intervals, and the contractile force was measured at an acquisition rate of 1 kHz. At 5 min after the 200 Hz stimulation, fatigue was measured under 50 repeated 500-ms, 100-Hz pulses at a 2-s interval. After muscle length was measured, the muscles were removed from the chamber, the tendons were dissected out, and muscle weight was measured. Specific force was calculated from the absolute force, muscle weight, and muscle length, assuming a density of 1.056 g/ml.

### 2.8. Histology

OCT embedded EDL samples and liver samples were mounted on a precooled platform and sectioned at 10 μm thickness with a cryostat (CM1510; Leica, Wetzlar, Germany) at −20°C and mounted on polylysine-coated slides. For EDL samples, whole sections were stained for hematoxylin-and-eosin (H&E) to assess cross-sectional area (CSA) or Sirius Red staining to determine fibrosis. Stained slides were imaged with a fully automated wide-field light microscope (Nikon Corp.) with a 20× objective lens. Images were captured using a high-sensitivity Andor Clara CCD camera. Sirius red staining was performed as previously described (34499764). Briefly, sections were fixed in Bouin’s fixative for 1 hour at 56°C, stained for 1 hour in Master Tech Picro Sirius Red, and washed in 0.5% glacial acetic acid before mounting with Permount. For liver samples, whole sections were stained for H&E and Oil Red staining to determine fat droplet accumulation.

### 2.9. Muscle imaging

FDB muscles were digested in KHB containing 2 mg ml^−1^ collagenase type II (Worthington Biochemical) for 2 h at 37 °C. Digested FDB muscles were transferred to KHB without collagenase or CaCl2 and titrated to separate them into individual muscle fibers. Fibers were plated on a laminin-coated 96-well plate. Fibers were used within 5 h of dissection. Fibers were loaded with 5 μM BODIPY C11 (D3861, Invitrogen), 2 μM DCFH-DA (R252, Dojin, Japan), 2 μM FerroOrange (F374, Dojin, Japan) and 10 μM Liperfluo (L248, Dojin, Japan) in the dark for 30 min (BODIPY C11, DCFH-DA, and FerroOrange) or 1 hr (Liperfluo) at room temperature (22-25 °C). Loaded fibers were imaged using a confocal microscope (STELLARIS5, Leica).

### 2.10. Intracellular Ca^2+^ measurement

The Intracellular Ca^2+^ measurement was assessed in intact FDB muscles, as previously described [24]. Briefly, the collagenated FDB muscles were plated on a laminin-coated 96-well plate. Single FDB fibers were loaded with 5 μM fluo-4 AM (Invitrogen), 5 μM Rhod-2 (Invitrogen) and Pluronic F-127 (P3000MP, Invitrogen) in the dark for 30 min at room temperature (22–25°C), followed by a washing with dye-free KHB. Fibers were then washed in dye-free solution supplemented with 25 μM N-benzyl-p-toluenesulfonamide (BTS, AAJ64910MA, Fisher Scientific) for 20 min to block myosin ATPase. To stimulate Ca^2+^ release from the sarcoplasmic reticulum (SR), fibers were injected with 2 mM 4-CMC, an agonist ryanodine receptor. The peak fluorescence intensity value was measured after 4-CMC stimulation, and the quantification data were plotted as changes from the pre-stimulation level.

### 2.11. Western blotting

Whole muscle lysates were utilized for western blotting. Approximately 20-30 mg of frozen GAS muscle was cut, weighed, and homogenized in a ground-glass homogenization tube using a mechanical ground glass pestle grinder in 10 volumes of ice-cold RIPA buffer (08714-04, Nacalai Tesque Inc) supplemented with protease phosphatase inhibitor cocktail. Homogenates were centrifuged for 15 minutes at 12,000 x g at 4°C prior to protein assessment of the supernatant via BCA (Thermo Scientific, 23227). Equal protein was then mixed with Laemmli sample buffer with BME to a final protein concentration of 2 mg/mL and denatured by incubating at 95° C for 5 minutes. 10 μg of protein was then loaded onto a 4-15% gradient TGX gel (Bio-Rad) and separated via electrophoresis. Proteins were transferred to nitrocellulose membranes on ice and then ponceau stain was placed on the membrane to visualize the membrane bound protein. Membranes were then blocked in Blocking One at room temperature with rotation and then incubated in primary antibodies targeting 4HNE (1:1000, Abcam, ab48506), ACSL4 (1:1000, Santa Cruz Biotechnology, sc-365230), LPCAT3 (1:1000, Proteintech, 67882-1-Ig), ALOX12 (1:1000, Santa Cruz Biotechnology, sc-365194), Gpx4 (1:1000, Abcam, ab125066) and GAPDH (1:5000, Merck Millipore, CB1001) in overnight at 4° C with gentle rocking. Membranes were then washed with TBST and incubated in species-appropriate 159 secondaries (1:5,000) in 3% milk in TBST for 1 hour at room temperature with rotation. Then, the membranes were incubated with the appropriate secondary antibody conjugated to horseradish peroxidase and visualized with SuperSignal reagent in a Bio-Rad ChemiDoc imaging system. Image J software was used for densiometric analysis of bands resolved at the predicted molecular weights.

### 2.12. Statistical analysis

Values are expressed as mean ± SEM. Statistical comparisons were performed using an unpaired two-tailed Students t-test for 2-group analyses, two-way ANOVA with Sidak’s post-hoc test for multiple comparisons (GraphPad Prism 8.1.0).

## 3. Results

### 3.1. D-PUFA does not change food/water intake and body weight from STZ-induced diabetic mice

For both males and females, food intake and water intake in the H-PUFA and D-PUFA cohort were not different (Fig. 1B-G). Following injection of STZ, both H-PUFA and D-PUFA groups lost weight similarly, and no difference in body weight between the groups was observed (Fig. 1J). Consistent with a previous study (20194573), STZ diabetic mice have lower grip strength compared with the vehicle groups, but higher grip strength was observed in D-PUFA groups (Fig. 1M).

### 3.2. There was a trend for D-PUFA protection against glucose intolerance from STZ-induced diabetic mice

Consistent with published work [25, 26], we confirmed that the fasting blood glucose levels and the area under the curve (AUC) for GTT were significantly greater in both male and female STZ mice than in vehicle groups (Fig. 2A-D), while D-PUFA tend to lower AUC of GTT in male mice (Fig. 2A). In addition, we evaluated the mass of white adipose tissue and liver (Fig. 2E-H). There were no differences in the mass of inguinal white adipose, epididymal white adipose tissue, and liver between H-PUFA and D-PUFA groups (Fig. 2E-G). It is noteworthy that D-PUFA did not develop hepatic steatosis in STZ-induced diabetic mice (Fig. 2H).

**Figure 2.**
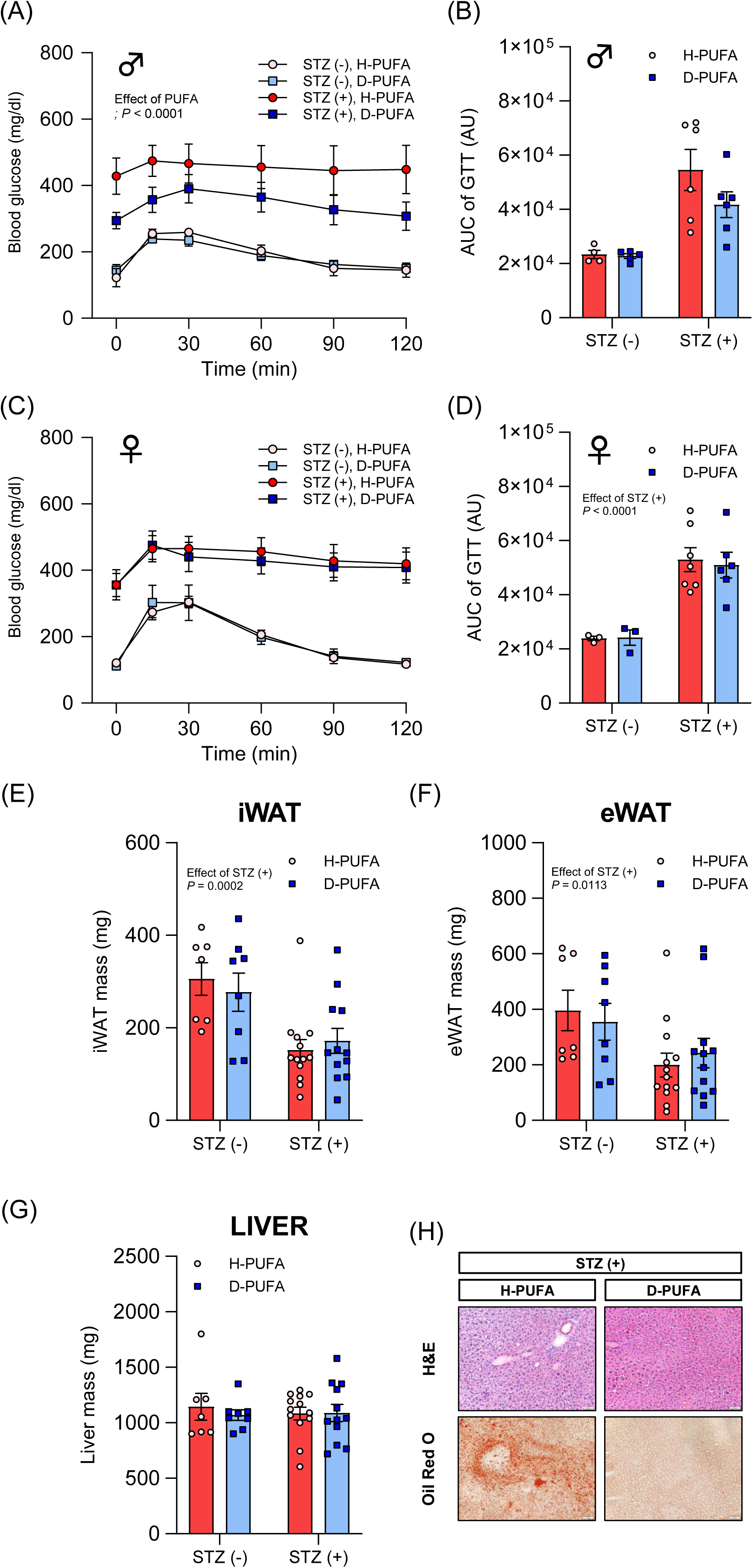
Characterization of D-PUFA for glucose intolerance from STZ-induced diabetes. (A and B) Intraperitoneal glucose tolerance test (A) and AUC (B) from glucose tolerance test in male mice (n = 4-6/group). (C and D) Intraperitoneal glucose tolerance test (C) and AUC (D) from glucose tolerance test in female mice (n = 3-7/group). (E) Inguinal white adipose tissue mass. (F) Epididymal white adipose tissue mass. (G) Liver mass. (H) Representative images of H&E and Oil Red O staining. For the group comparison, two-way ANOVA were performed with Sidak’s post-hoc test. All data are mean ± SEM.

### 3.3. D-PUFA reduce lipid peroxidation in skeletal muscle from STZ-induced diabetic mice

To assess the lipid peroxidation and oxidative status of STZ-induced diabetic mice, we conducted an imaging analysis from confocal microscopy. Compared to the vehicle group, the STZ mice showed significantly higher oxidized BODIPY fluorescence intensity from H-PUFA, while lower fluorescence intensity was observed in D-PUFA groups (Fig. 3A and B). Similar to BODIPY results, D-PUFA had a decreased DCFH and Liperfluo intensity from STZ-induced diabetic muscle (Fig. 3C-E), without a decrease in FerroOrange intensity to assess the iron accumulation (Fig. 3F). Indeed, D-PUFA were sufficient to suppress the accumulation of 4-hydroxynonenal (4-HNE) from STZ-induced diabetic muscle (Fig. 3G and H). These data indicate that D-PUFA potently reduces lipid peroxidation of skeletal muscle tissue induced by hyperglycemia.

**Figure 3.**
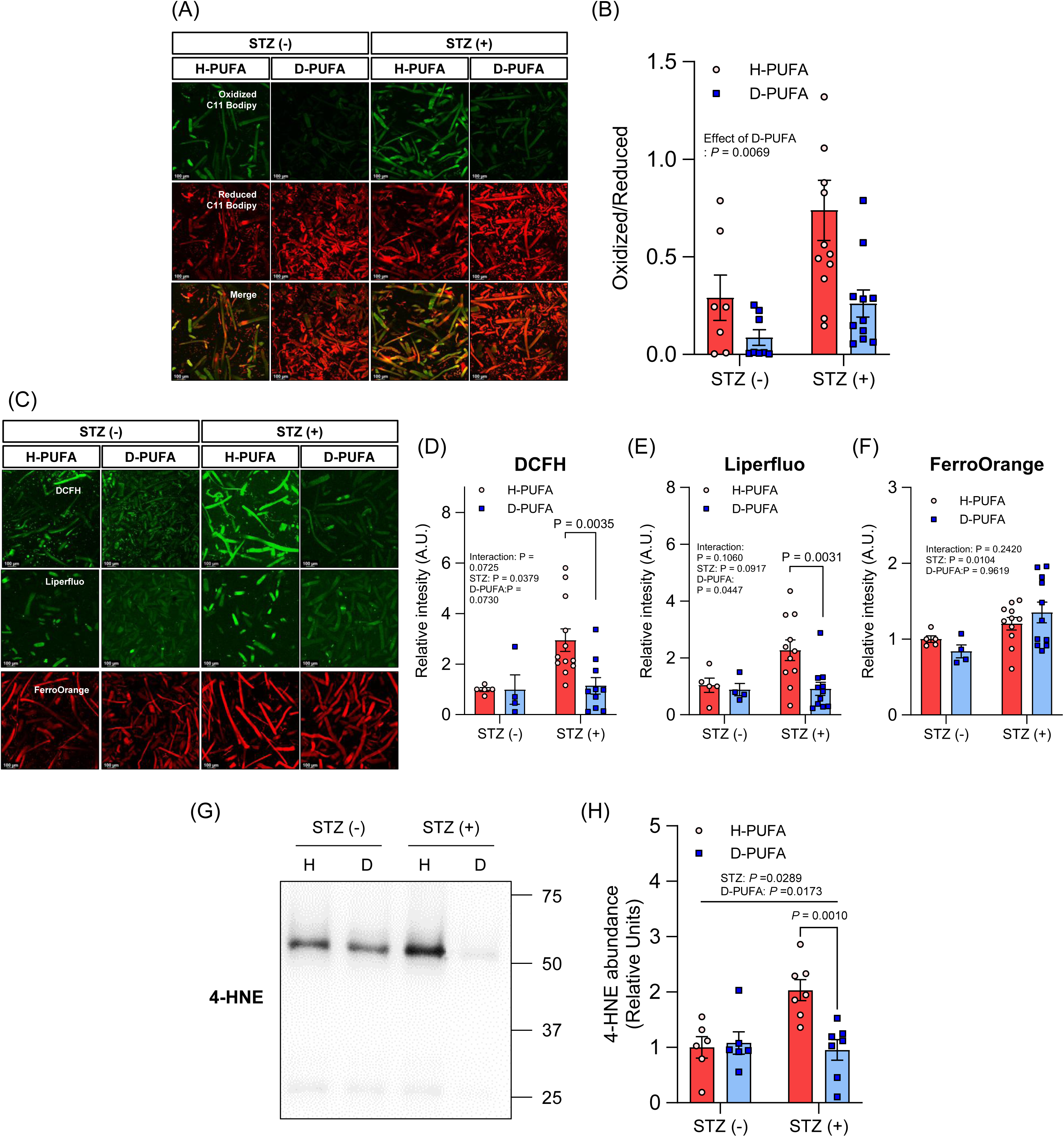
D-PUFA protects lipid peroxidation in skeletal muscle from STZ-induced diabetes. (A and B) Representative images of confocal microscopy (A) and quantification of fluorescence intensity (B) for BODIPY C11 corresponding to isolated fibers from the FDB muscle (n = 7-12/group). (C-F) Representative images of confocal microscopy (C) and quantification of fluorescence intensity for DCFH (D), Liperfluo (E), and FerroOrange (F) corresponding to isolated fibers from the FDB muscle (n = 4-12/group). (G and H) Representative images (G) and quantification (H) of protein abundance for 4-hydroxynonenal (4-HNE) (n = 6-7/group). For the group comparison, two-way ANOVA were performed with Sidak’s post-hoc test. All data are mean ± SEM.

### 3.4. D-PUFA protects muscle atrophy and fibrosis from STZ-induced diabetic mice

D-PUFA had no significant effect on mass of hindlimb muscle in the vehicle groups (Fig. 4A), while D-PUFA was associated with higher muscle mass compared to H-PUFA in STZ groups (Fig. 4B). Indeed, CSA of EDL muscle was higher in D-PUFA compared to H-PUFA in STZ groups (Fig. 4C and D). These observations show that D-PUFA protects against loss of muscle mass even at the microscopic level. In addition, D-PUFA cohort showed decreased Sirius Red staining compared to H-PUFA animals, (Fig. 4C and E), suggesting that D-PUFA protects muscle fibrosis in hyperglycemia. Taken together, D-PUFA ameliorates atrophy and fibrosis in hindlimb muscle induced by hyperglycemia.

**Figure 4.**
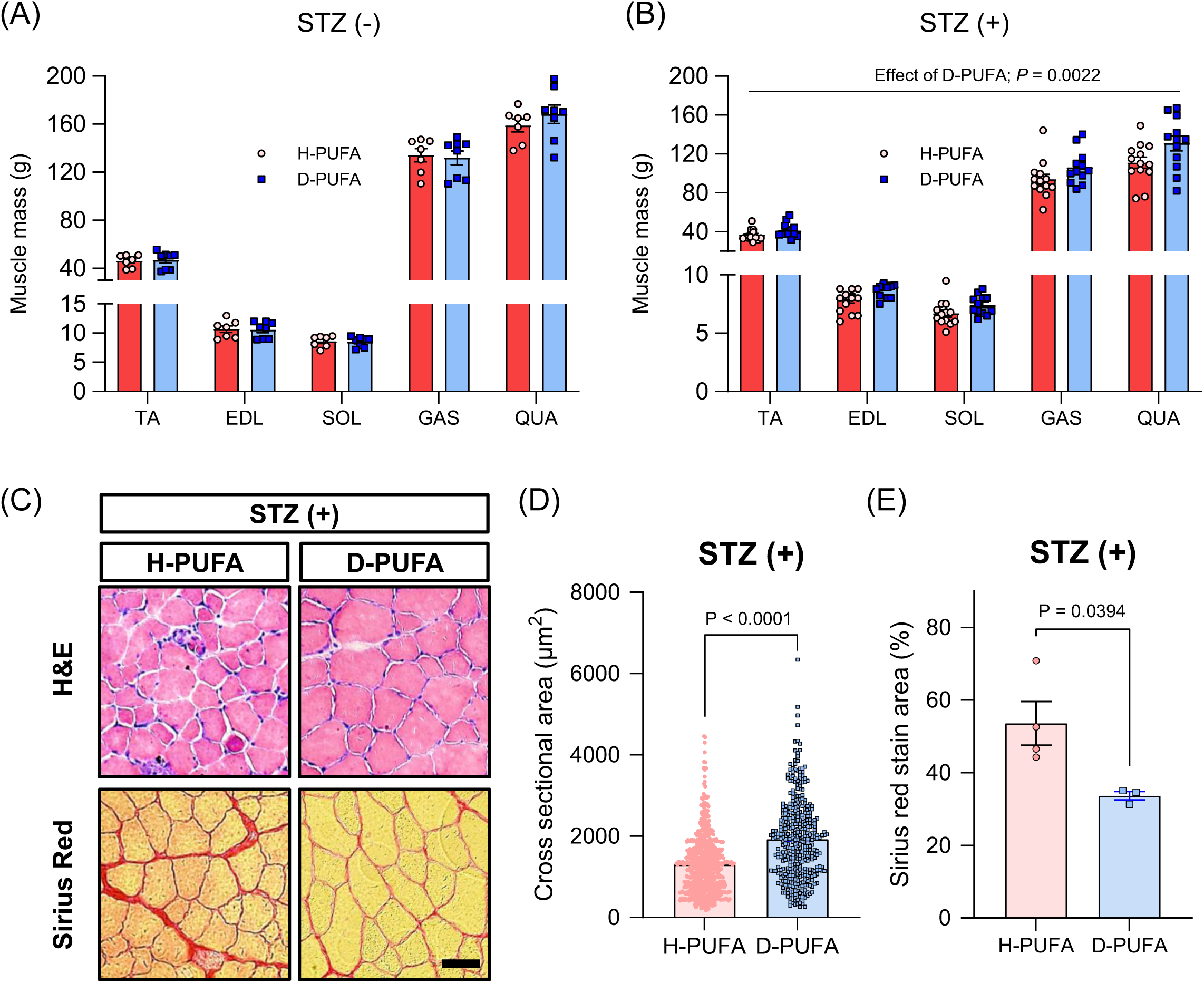
D-PUFA protects muscle atrophy from STZ-induced diabetes. (A and B) Hindlimb muscle mass from Vehicle (A) and streptozotocin injected (B). (C) Representative images of H&E and Picro-Sirius Red staining. (D) Quantification of H&E of any individual fibers (n = >695). (E) Quantification of Picro-Sirius Red staining (n = 3-4/group). Scale bar, 50 μm. For the multiple muscle mass comparison, two-way ANOVA were performed with Sidak’s post-hoc test. For the two-group comparison analysis, an unpaired two-tailed t-test were performed. All data are mean ± SEM.

### 3.5. D-PUFA protects muscle weakness from STZ-induced diabetic mice

The force-frequency relationship and fatigue resistance were analyzed for EDL muscle with the vehicle (Fig. 5A-D) and STZ groups (Fig. 5E-H). In the vehicle groups, twitch and tetanic muscle force production were not different between H-PUFA and D-PUFA. As expected, STZ significantly reduced twitch and tetanic force production in H-PUFA, while D-PUFA cohort had higher force production (Fig. 5E-G). No differences in fatigue resistance were found by repeated tetanic contractions between H-PUFA and D-PUFA in both vehicle and STZ groups (Fig. 5D and H). Our published study demonstrated that muscle weakness in diabetes is associated with SR-mediated impairment of the [Ca^2+^]i flux [19, 27]. Therefore, we examined the effect of D-PUFA on [Ca^2+^]i flux using a direct activator of RyR, 4-CmC [23]. For Fluo-4 (Fig. 5J) and Rhod-2 (Fig. 5K) fluorescence intensity, D-PUFA had higher Ca^2+^ peak levels following 4-CmC administration compared to H-PUFA in STZ groups (Fig. 5I-K), suggesting that D-PUFA protects muscle weakness through mechanisms involving calcium homeostasis.

**Figure 5.**
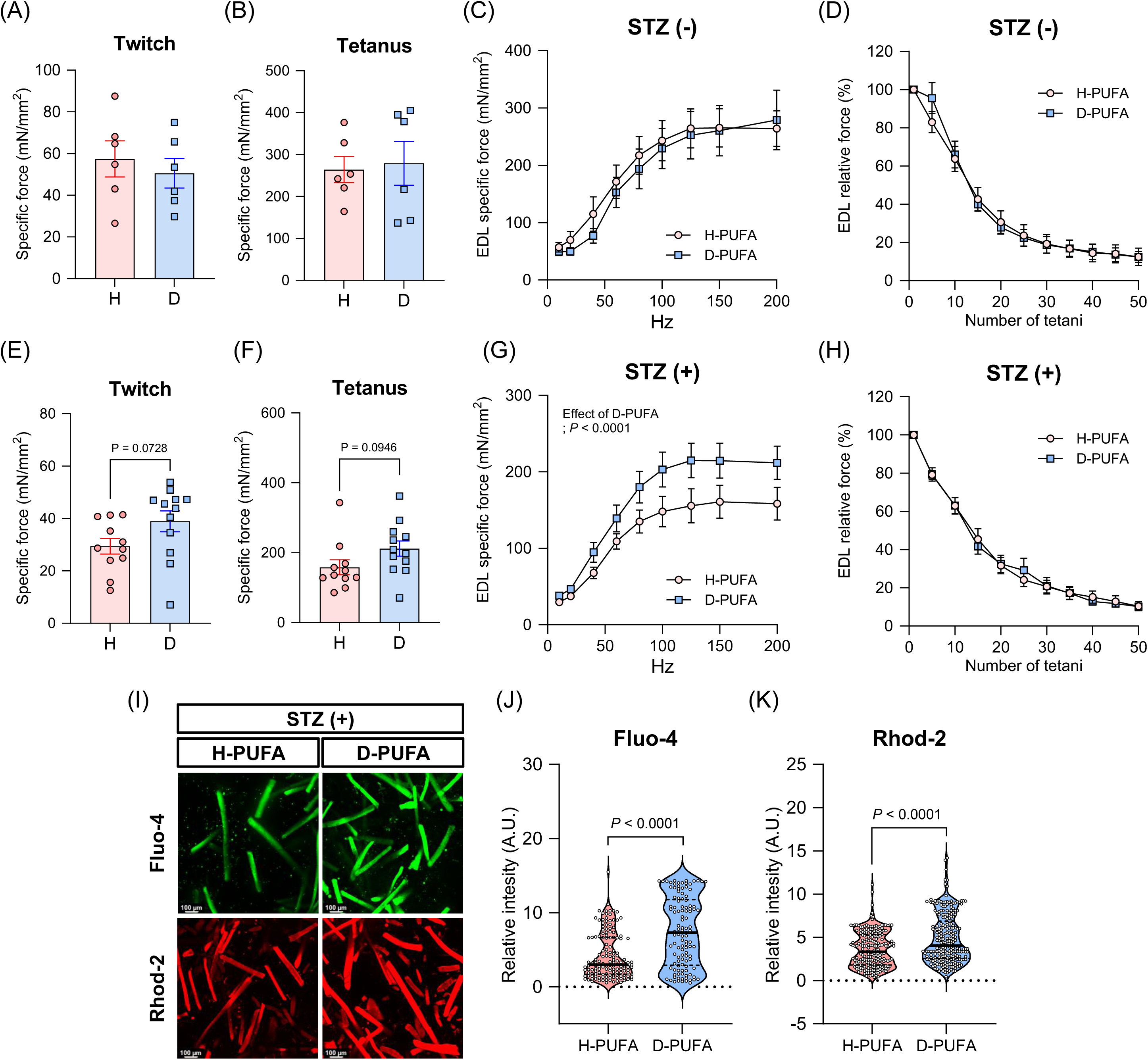
D-PUFA protects muscle weakness from STZ-induced diabetes. (A-D) Contractile properties in extensor digitorum longus (EDL) muscles of vehicle. Peak twitch forces (A), maximal tetanic forces (B), force-frequency curve (C) and fatigue resistance with repeated tetanic stimulation (D) (n = 6/group). (E-H) Contractile properties in EDL muscles of streptozotocin injected. Peak twitch forces (E), maximal tetanic forces (F), force-frequency curve (G) and fatigue resistance with repeated tetanic stimulation (H) (n = 11-12/group). (I-K) Calcium imaging. (I) Representative immunofluorescence images of Fluo-4 and Rhod-2 in FDB muscle fibers. Quantification of Fluo-4 (J) and Rhod-2 (K) fluorescence intensity in 2 mM 4-CMC-stimulated FDB fibers (n = >695). For the force-frequency curve and fatigue resistance experiments, two-way ANOVA were performed with Sidak’s post-hoc test. For the two-group comparison analysis, an unpaired two-tailed *t*-test were performed. All data are mean ± SEM.

### 3.6. D-PUFA does not alter Ferroptosis-related protein abundance

We investigated the abundance of ferroptosis-associated proteins such as ACSL4, LPCAT3, ALOX12, and Gpx4 to identify the molecule mechanisms (Fig. 6A). No significant differences were observed in these proteins which were comparable among the groups (Fig. 6B-F).

**Figure 6.**
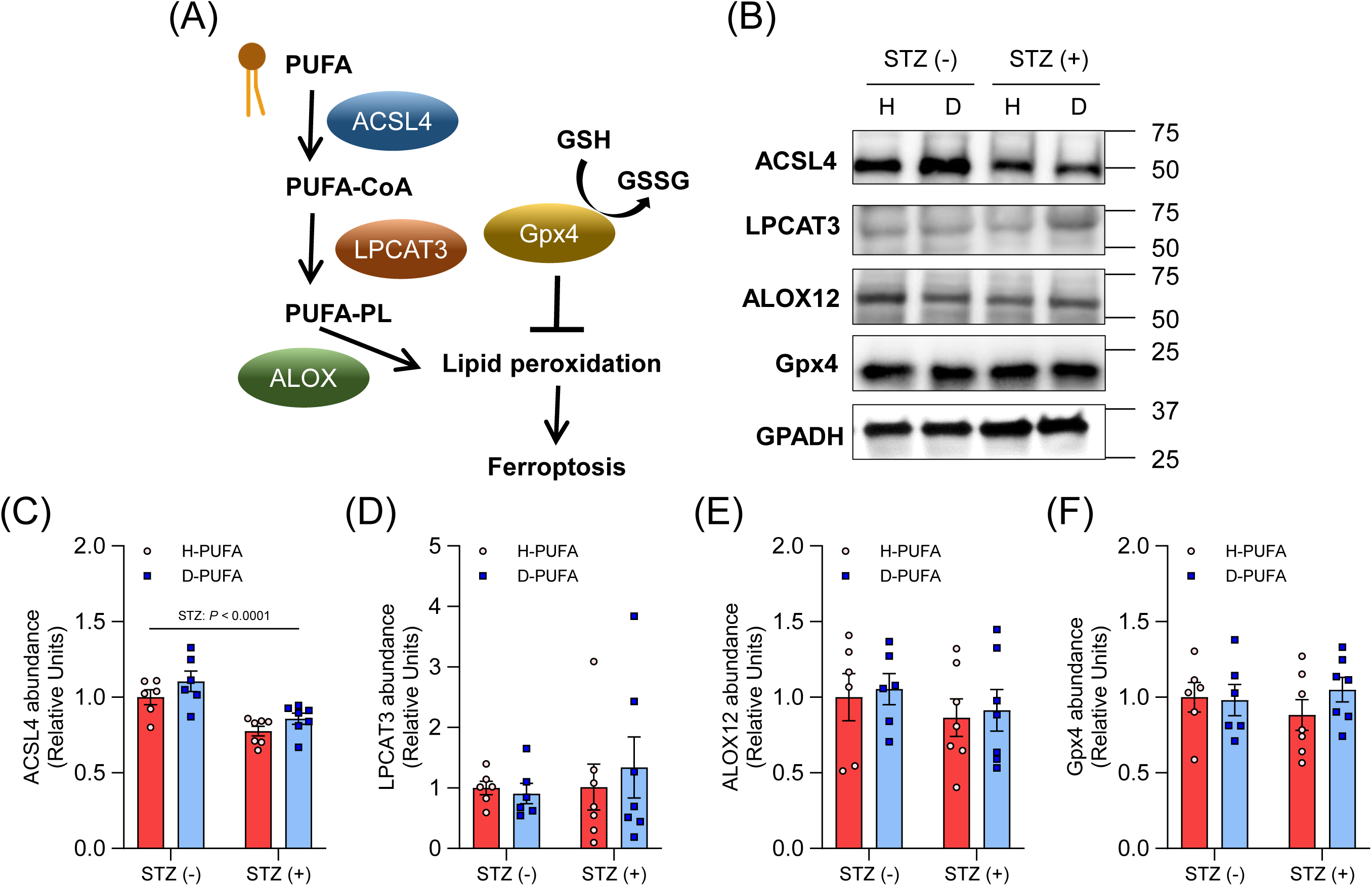
D-PUFA does not alter Ferroptosis-related protein abundance. (A) Schematic of Ferroptosis pathway. (B) Representative images of protein abundance. Quantification of protein abundance for ACSL4 (C), LPCAT3 (D), ALOX12 (E) and Gpx4 (F) (n = 6-7/group). For the group comparison, two-way ANOVA were performed with Sidak’s post-hoc test. All data are mean ± SEM.

## 4. Discussion

Oxidative stress has been implicated in diabetic myopathy [5]. A previous study demonstrated that antioxidants could attenuate muscle atrophy and weakness in type 1 diabetes, suggesting that ROS may directly induce muscle atrophy [28]. However, antioxidants may change redox activity to other cellular systems resulting in protection against loss of muscle mass and weakness. We recently discovered that suppression of lipid peroxidation and their reactive lipid aldehydes is sufficient to prevent disuse-induced muscle atrophy [8]. Therefore, this study uses a D-PUFA-based suppression of lipid peroxidation to more definitively conclude its role in mediating diabetic myopathy. Importantly, suppression of lipid peroxidation by D-PUFA protects against muscle atrophy and weakness induced by diabetes, suggesting that accumulated lipid peroxidation in skeletal muscle is mainly the mechanism by which muscle atrophy and weakness are induced by diabetes.

D-PUFA did not affect muscle mass of healthy mice while D-PUFA had significantly higher muscle mass than H-PUFA in STZ-induced diabetic mice. Recently, we described lipid peroxidation as a factor in skeletal muscle atrophy with age and disuse [7, 8]. The oxidation of membrane lipids destroys lipid bilayer packing causing lipid hydroperoxide accumulation.

Indeed, oxidation of phosphatidylethanolamine leads to a generation of ferroptosis [29]. A previous study showed that liproxstatin-1, an inhibitor of LOOH propagation, protects against muscle atrophy with denervation [30]. Accordingly, we believe lipid hydroperoxide likely contributes to muscle atrophy induced by several states including disuse, aging, diabetes, and denervation.

Many studies have shown that diabetes causes impairment of muscle strength and power [31, 32]. However, the mechanisms underlying muscle weakness in diabetes mellitus have not yet been elucidated. Skeletal muscle contraction is determined via a Ca^2+^-dependent process known as excitation-contraction coupling [33]. We recently demonstrated that the impairment of Ca^2+^ release from the SR has been linked to muscle weakness in diabetes [19, 23]. A previous study demonstrated that ryanodine receptor was oxidized and Ca^2+^ leak from SR contributed to muscle weakness in aged mice [34], suggesting that diabetic muscle weakness may be associated with oxidation of ryanodine receptors in skeletal muscle. Indeed, a recent study demonstrated that the endoplasmic reticulum membrane is a key site of lipid peroxidation [35]. Accordingly, our data shows that D-PUFA protects against muscle weakness depending on the improvement of decreased intracellular Ca^2+^ release levels (Fig. 5I-K).

Recently, our [8] and other laboratories [11, 36, 37] demonstrated that lipid aldehyde contributes to muscle atrophy and weakness. Several studies show that overexpressing Gpx4 protects against muscle atrophy with suppression of lipid peroxidation in aging [11, 37] and denervation [30]. Another study demonstrated that Gpx4 haploinsufficiency accelerates cardio-metabolic derangements induced by obesity [38], suggesting that suppression of lipid aldehyde is a countermeasure to metabolic disease. Together, our study is the first to demonstrate that D-PUFA can directly suppress lipid peroxidation, protecting against muscle atrophy and weakness in diabetes, without any observed side-effects.

The iron-dependent lipid peroxidation is the hallmark of ferroptosis-induced cell death [39]. Ferroptosis has been linked to several pathophysiology such as neurodegenerative disorders, cancers, and kidney diseases [40, 41]. To gain insight into the role of lipid peroxidation induces muscle atrophy and weakness induced by diabetes, we evaluated the abundance of ferroptosis-related proteins. Arachidonic acid-mediated initiation of ferroptosis depends on an acyl-coenzyme A (CoA) synthetase long-chain family member 4 (ACSL4) and then it upregulates LPCAT3 [29, 42]. Lipoxygenase (LOX) generates doubly and triply-oxygenated-diacylated phospholipid species and acts as a signal for ferric ions and, ultimately, promotes ferroptosis [29]. However, there were no differences in these protein abundances in any group (Fig. 6 A-D). Therefore, ACSL4/LPCAT3/15-LOX pathway may not be important in the production of lipid peroxidation with diabetic myopathy.

The present study suggests that D-PUFA protects muscle atrophy and weakness induced by diabetes independent of ACSL4/LPCAT3/15-LOX pathway. Gpx4 acts as a master regulator in the ferroptosis process [43], but D-PUFA did not alter the abundance of Gpx4 protein in STZ-induced diabetic mice. Another mechanism may potentially be the regulation of intracellular organelles such as mitochondria. In cardiac muscle, STZ-induced diabetic mice decreased cardiolipin content while cardiolipin transgenic mice attenuated maladaptive cardiolipin remodeling and bioenergetic inefficiency mitochondria induced by STZ treatment [44]. Indeed, muscle biopsies from type 2 diabetic patients tend to reduce cardiolipin content compared to nondiabetic lean subjects [45]. The cardiolipin substitutes for the oxidized one if oxidative stress is high in mitochondria. D-PUFA may decrease cardiolipin to get oxidized, resulting in attenuated muscle atrophy similar to a transgenic model. Furthermore, a previous study demonstrated that the product of dihydroorotate dehydrogenase (DHODH) in mitochondria attenuates ferroptosis induced by inhibition of Gpx4 [46]. However, the role of DHODH in diabetic myopathy is still unclear. Inflammation is a well-known factor in diabetes-related muscle degeneration [4]. Multiple mechanisms can be behind the degeneration, such as an impairment of protein synthesis in muscles through pro-inflammatory cytokine-activated STAT3 [47]. Accordingly, the protection we describe may have to do with the anti-inflammatory properties of D-PUFAs, which down-regulate inflammation by decreasing the levels of pro-inflammatory eicosanoids [48], as demonstrated the extent of the protection in a murine lung inflammation model [49, 50]. Together, the mechanisms of D-PUFA protect muscle atrophy and weakness in diabetes remain unknown, and further investigations are required to clarify.

## 5. Conclusions

We examined the effect of deuterated PUFA on muscle atrophy and weakness in STZ-induced diabetic mice. Our findings indicate that D-PUFA protects against muscle atrophy induced by type 1 diabetes. Additionally, we found that preventing muscle weakness by D-PUFA may be explained by intracellular calcium release from SR in diabetes. We also found that the prevention of muscle atrophy by D-PUFA is not mediated by ALOX/ACSL/LPCAT3 and Gpx4 pathways. Our finding suggested that D-PUFA may be a promising therapeutic approach for type 1 diabetes-induced muscle atrophy and weakness in skeletal muscle.

## Acknowledgements

We thank Undergraduates students for cross sectional are analysis support.

